# Modeling glioblastoma tumor progression via CRISPR-engineered brain organoids

**DOI:** 10.1101/2024.08.02.606387

**Authors:** Matthew Ishahak, Rowland H. Han, Devi Annamalai, Timothy Woodiwiss, Colin McCornack, Ryan T. Cleary, Patrick A. DeSouza, Xuan Qu, Sonika Dahiya, Albert H. Kim, Jeffrey R. Millman

## Abstract

Glioblastoma (GBM) is an aggressive form of brain cancer that is highly resistant to therapy due to significant intra-tumoral heterogeneity. The lack of robust in vitro models to study early tumor progression has hindered the development of effective therapies. Here, we develop engineered GBM organoids (eGBOs) harboring GBM subtype-specific oncogenic mutations to investigate the underlying transcriptional regulation of tumor progression. Single-cell and spatial transcriptomic analyses revealed that these mutations disrupt normal neurodevelopment gene regulatory networks resulting in changes in cellular composition and spatial organization. Upon xenotransplantation into immunodeficient mice, eGBOs form tumors that recapitulate the transcriptional and spatial landscape of human GBM samples. Integrative single-cell trajectory analysis of both eGBO-derived tumor cells and patient GBM samples revealed the dynamic gene expression changes in developmental cell states underlying tumor progression. This analysis of eGBOs provides an important validation of engineered cancer organoid models and demonstrates their utility as a model of GBM tumorigenesis for future preclinical development of therapeutics.

## MAIN

Mapping and understanding the genetic drivers of tumorigenesis and tumor progression have been fundamental goals in the study of cancer. The Cancer Genome Atlas (TCGA), which was started in 2006, has performed molecular characterization of over 20,000 primary tumors from thirty-three cancer types^1^. Glioblastoma (GBM), the first cancer studied by TCGA^2^, is the most common primary malignant brain tumor in adults, accounting for over 14% of all brain and other central nervous system tumors in the United States^3^. Patients with GBM have a poor prognosis, with a median survival of less than two years after diagnosis despite rigorous therapy^4^. To date, TCGA has conducted comprehensive multidimensional analysis of over six hundred GBM tumors, resulting in the classification of GBM into Proneural, Neural, Classical, and Mesenchymal subtypes^5^. Despite thorough characterization of patient samples, very few new therapeutic options for GBM have been developed in last the decade^6^, which highlights the need for improved pre-clinical models.

Currently, the most widely used pre-clinical models for brain tumors include genetically modified mice, syngeneic mouse models, and patient-derived cancer spheroids^7-9^. Genetic modification of tumor suppressor genes and introduction of oncogenic perturbations in mouse models result in the formation of GBM-like tumors. However, species-specific differences in brain morphology and physiology combined with the differences in tumor growth patterns and antigen expression limit the human relevance of mouse models. While patient-derived three-dimensional (3D) tumor spheroids can recreate the cellular heterogeneity of GBM, these models are often derived from late-stage tumors which hinder investigation of early tumorigenesis processes due to the high mutational burden that occurs over time^10,11^.

Recently, human pluripotent stem cell-derived brain organoids have emerged as a powerful in vitro system to model human neural development and diseases^12-15^. Brain organoids are of particular interest for modeling cerebral tumors, since it has been observed that tumor cell types are more closely associated with developmental-like signatures, rather than adult cell populations^16^ and that neural development transcriptional programs govern invasion^17,18^. Previously, CRISPR-Cas9-based methods have been utilized to introduce oncogenic mutations into developing organoids resulting in neoplastic cerebral organoids^8,19,20^. While these model systems exhibit many features of cancer, it remains unclear how well they recapitulate human GBM tumorigenesis and if the resulting tumors are comparable to advanced GBM patient tumors. As a result, there is a need for further development and characterization of engineered organoid models of GBM.

Here, we generated engineered GBM brain organoids (eGBOs) from genetically modified stem cells harboring mutations of both the Proneural and Mesenchymal subtypes of GBM. Using single-cell RNA sequencing (scRNAseq) and spatial transcriptomic analyses, we demonstrate that GBM-associated mutations alter neural development in brain organoids. Further, we find that transplantation of cells from eGBOs into mice results in tumors that recapitulate subtype-specific characteristics of human GBM tumors. Finally, we identify dynamic gene expression changes in developmental cell states underlying tumor progression using integrated single-cell transcriptomic data from eGBOs and patient GBM tumors. These eGBOs are an effective model of GBM tumorigenesis and, combined with the robust framework for computational analysis we describe, provide an important validation of engineered cancer organoid models for future development of cancer therapies.

## RESULTS

### Generation of eGBOs from stem cells harboring GBM-associated mutations

Prior characterization of the transcriptomes from human GBM tumors identified genetic subtypes of GBM with specific mutations associated with different subtypes^5^. Therefore, we first sought to genetically engineer stem cells to mimic the heterogeneous genetic landscape of GBM. Through serial introduction of GBM-associated mutations in healthy human pluripotent stem cells (hPSCs) using CRISPR-Cas9, we developed two engineered GBM stem cell (eGSC) lines harboring representative mutations of the Proneural (PRO) and Mesenchymal (MES) GBM subtypes (Fig. 1a, Extended Data Fig. 1a). First, hPSCs were transfected with a CRISPR-Cas9 ribonucleoprotein (RNP) complex to introduce the machinery necessary for precise genetic modification. We then introduced a C228T mutation in the telomerase reverse transcriptase (*TERT*) promoter region to create a genetic background that promotes rapid tumor progression through reactivation of the *TERT* gene, a characteristic of GBM progression driven by non-canonical NF-κB signaling^10,21,22^.

**Fig. 1:**
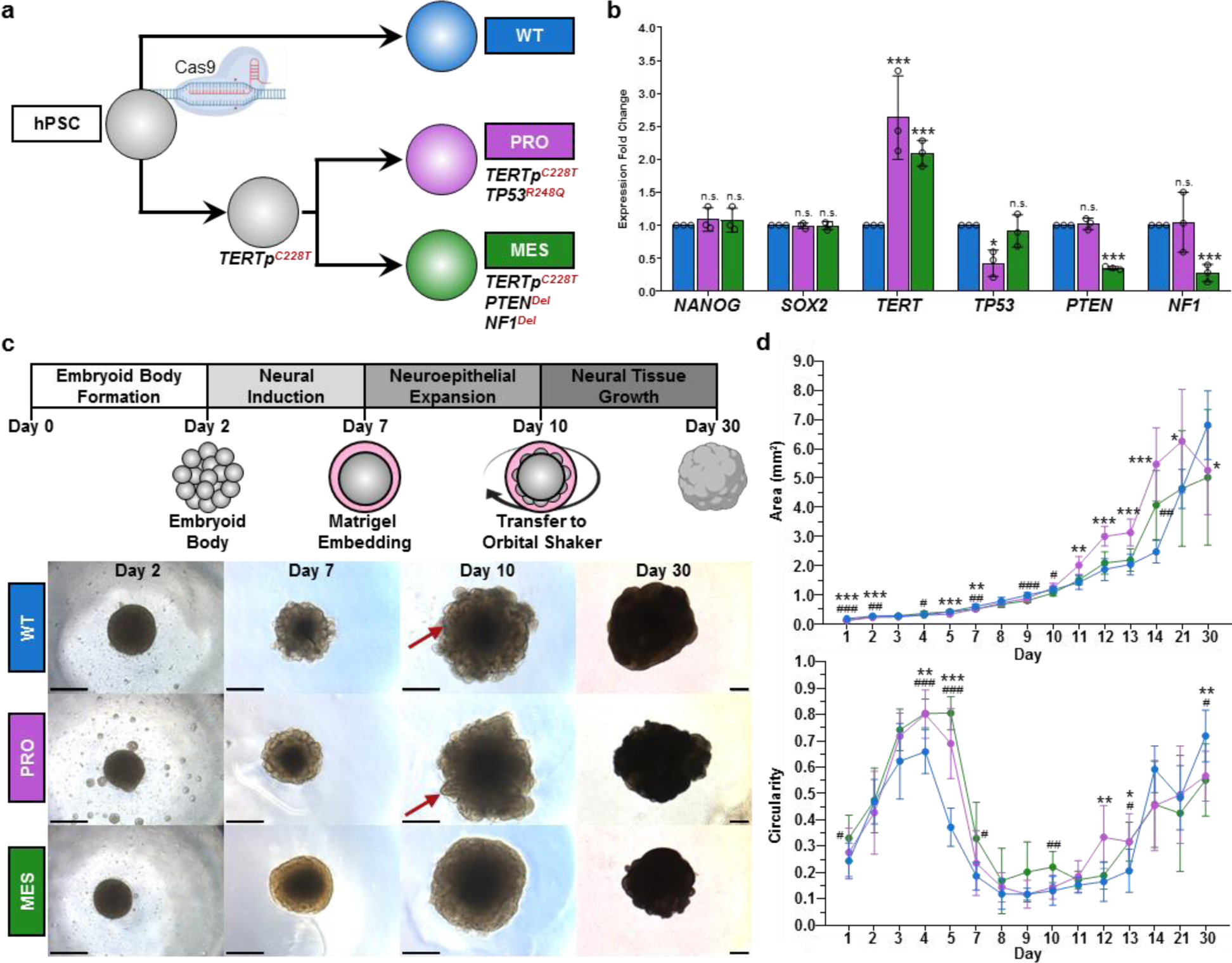
Generation of eGBOs from hESCs harboring GBM-associated mutations. **a,** Genetic engineering strategy to generate eGSCs by introducing GBM-associated mutations into hPSCs via sequential CRISPR-Cas9 gene editing. **b,** qPCR analysis of genetically engineered hPSCs indicating unchanged expression of pluripotency markers and altered expression of genes targeted by CRISPR editing. Statistical analysis by parametric unpaired t-test versus WT condition with Holm-Sidak correction for multiple comparisons. *P<0.05; **P< 0.01; ***P<0.0001 **c,** Schematic of brain organoid differentiation protocol and brightfield images during cerebral organoid differentiation. Red arrowheads indicate neuroepithelial growth, which is slight reduced in PRO organoids, and greatly reduced in MES organoids. Scale bars = 500µm **d,** Surface area (top) and circularity (bottom) measurements during organoid differentiation (n = 10). Statistical analysis by two-way ANOVA using Geisser-Greenhouse correction and Tukey correction for multiple comparison (* indicates WT vs PRO comparisons and # indicates WT vs. MES comparisons). *P<0.05; **P< 0.01; ***P<0.0001.

The p53 pathway is dysregulated in approximately 85% of all GBM tumors^23^ and has been implicated in invasion, migration, proliferation, evasion of apoptosis, and cancer cell stemness^24-28^. Despite being the most frequently gene mutated in GBM, *TP53* mutations are mostly commonly observed in the PRO subtype of GBM^5^. Therefore, we introduced a R248Q hotspot mutation in the *TP53* gene to recapitulate the genetic profile of the PRO GBM subtype.

Decreased expression of *NF1* is characteristic of the MES subtype of GBM^5,29^, caused by loss of the chromosome region containing the *NF1* gene. Additionally, the *PTEN* gene has been identified as a tumor suppressor that is recurrently mutated in GBM^30^, with deletion or mutation leading to dysregulation of the PI3K pathway^31^. Concomitant mutation of *NF1* and *PTEN* is found almost exclusively within MES subtype tumors^5^, therefore small deletions were introduced to the *NF1* and *PTEN* genes to recreate the genetic profile of the MES GBM subtype.

After introducing GBM-associated mutations, we validated that eGSCs maintained normal stem cell morphology and pluripotency marker expression. Quantitative PCR analysis confirmed that CRISPR genetic modifications significantly altered target gene expression, without altering expression of pluripotency markers (Fig. 1b). A two-fold increase in *TERT* expression was observed in both the PRO and MES eGSCs compared to wildtype (WT) hPSCs. Additionally, expression of *TP53* was significantly decreased in the PRO eGSCs, while both *NF1* and *PTEN* expression were significantly decreased in the MES eGSCs. Further, there was no significant difference in the expression of the stem cell marker genes, *NANOG* and *SOX2*. Both PRO and MES eGSCs also maintained characteristic stem cell morphology and expression of NANOG protein (Extended Data Fig. 1b). Next-generation sequencing (NGS) further confirmed highly efficient modifications in the *TP53*, *PTEN*, and *NF1* genes, and the *TERT* promoter region (Extended Data Fig. 1c). These results demonstrate that GBM subtype-specific mutations can be introduced into hPSCs without altering stemness.

Previous reports have proposed that GBM hijacks normal neural development mechanisms to drive tumor growth^17,18^. Therefore, we differentiated eGSCs into brain organoids, following an established protocol^14,32^ with minor modifications, to investigate how GBM-associated mutations may alter neural development (Fig. 1c, Supplementary Table 1). Visual assessment of developing organoids by bright-field imaging revealed slight morphological differences between WT organoids and eGBOs (Fig. 1c). Notably, MES eGBOs lacked neuroepithelial growth on day 10. This observation is consistent with previous reports that disruption of the *PTEN* gene leads to delayed differentiation and altered surface folding in cerebral organoids^33^. We also performed assessment of surface area and circularity to quantitatively assess developing organoids (Fig. 1d). During the initial days of differentiation, WT organoids and eGBOs were approximately the same size. Around day 6, however, developing eGBOs were significantly smaller than WT organoids. This corresponded with eGBOs maintaining a highly circular morphology during the transition from embryoid body formation to the neural induction stage and, while developing PRO eGBOs were not significantly different from developing WT organoids on day 10, developing MES eGBOs maintained this circular morphology. Despite these differences in growth dynamics, our data demonstrates that brain organoids were successfully generated from both WT hPSCs and eGSCs harboring GBM subtype-specific mutations.

### Transcriptional landscape of neural development is altered in eGBOs

Differentiation of hPSCs into brain organoids results in multiple cell types representing the developing cerebral cortex, ventral telencephalon, retina, and other regions of neural development. Therefore, we performed scRNAseq to characterize the cellular composition and transcriptional profile of WT organoids and eGBOs. We implemented a cell hashing approach to sequence all three genotypes in a single, multiplexed batch, thereby reducing technical variability (Extended Data Fig. 2a). A total of 9,310 single cells were sequenced at an average read depth of 58,696 reads per cell and a median of 2,857 genes per cell. Cells with less than two hundred genes detected and genes detected in less than three cells were excluded from downstream analysis. We achieved a hashing efficiency of approximately 77% with 7,155 cells identified by a single hashtag, while 1,062 cells and 1,085 were identified as negative and doublets, respectively. After filtering out negative and doublet cells, expression of a single hashtag enabled clear delineation of cells from WT, PRO, and MES organoids (Extended Data Fig. 2b). Further quality control filtering to remove low-quality cells based on the percentage of mitochondrial genes resulted in 5,073 high-quality cells for further analysis. Pseudobulk analysis of GBM-associated genes we targeted for mutation further validated our hashing methodology, with increased *TERT* expression observed in eGBOs compared to WT organoids and reduced expression of *TP53* in PRO eGBOs and both *PTEN* and *NF1* in MES eGBOs (Extended Data Fig. 2c).

Unsupervised, hierarchical clustering and embedding into a two-dimensional (2D) space with uniform manifold approximation and projection (UMAP) enabled annotation of cell types and revealed differential clustering of cells from WT organoids and eGBOs (Fig. 2a). We identified sixteen distinct cell populations using the Louvain clustering algorithm. The gene expression profiles of these clusters were highly correlated to the pallium and midbrain regions of the developing brain, regardless of the cluster consisting of cells from WT organoids or eGBOs (Fig 2b, Extended Data Fig. 2d). Cell types within WT organoids and eGBOs were annotated by a combination of supervised assessment of marker gene expression and unsupervised comparison to reference datasets (Fig. 2c, Extended Data Fig. 2e-f). We observed clearly defined cell populations of normal brain organoid development, including radial glial cells (expressing *SOX2* and *NES*), neuronal cells (expressing *GAP43* and *DCX*), and mesenchymal cells (expressing *COL1A2* and *DCN*). We also identified off-target populations commonly found in brain organoids, including neural crest cells (expressing *SOX10* and *FOXD3*), retinal progenitors (expressing *SIX6*), microglia (expressing *C1QC*), and endothelial cells (expressing *PECAM1*). Notably, these off-target populations primarily consisted of cells derived from both PRO and MES eGBOs, while the majority of the neuronal population consisted of cells derived from WT organoids (Fig. 2d).

**Fig. 2:**
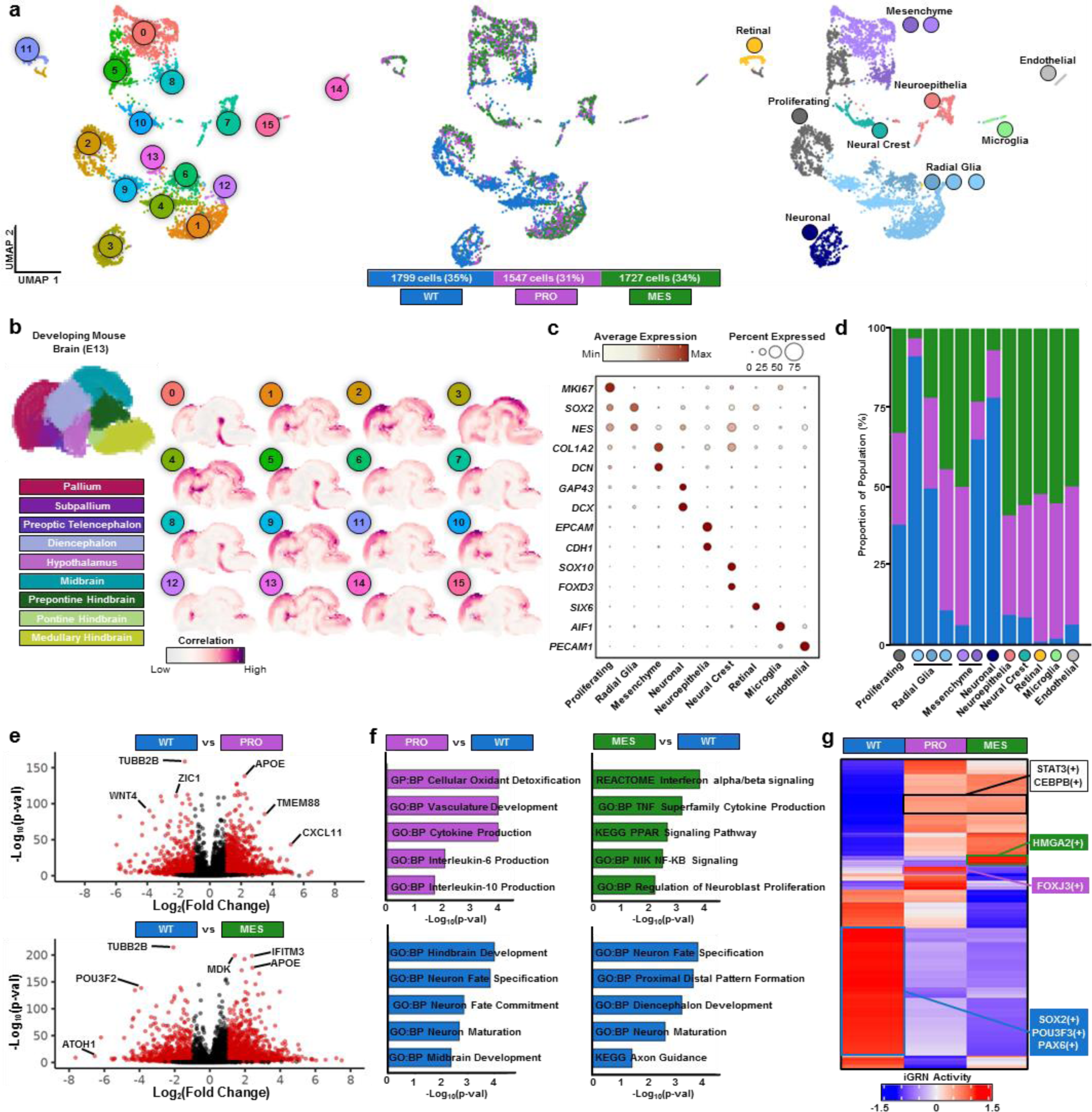
GBM mutations alter neural development gene regulatory networks in eGBOs. **a,** UMAPs showing Louvain clusters (left), sample identification (middle), and cell types (right) in brain organoids sequenced after 1 month of in vitro differentiation (5,073 total cells from WT, PRO and MES organoids). **b,** Spatial location inference of Louvain clusters identified in brain organoids. Reference regions provided by the Developing Mouse Brain database of Allen Brain Atlas (left) and correlation patterns of average regional marker gene expression for each Louvain cluster (right). **c,** Dot plot indicating expression of marker genes that define cell types observed in organoids. **d,** Stacked bar plot showing the proportion of cells from WT organoids and eGBOs for each cell type. **e,** Volcano plots for indicated pairwise comparisons from scRNAseq data. Red dots represent significantly differentially expressed transcripts (P<0.05) with Fold Change>=1. **f,** GSEA pathways from differentially expressed transcripts for indicated pairwise comparisons. **g,** Heatmap indicating scaled activity of iGRNs in WT, PRO, and MES organoids.

Next, we identified differentially expressed genes between WT organoids and eGBOs (Supplementary Table 2). We observed common neural development genes upregulated in WT organoids compared to eGBOs, (Fig. 2e). Specifically, *TUBB2B*, which encodes β-tubulin found in neuronal axons^34^, *ZIC1*, which regulates forebrain development^35^, *WNT4*, which regulates axonal growth^36^, *POU3F2*, which is involved in cortical development^37^, and *ATOH2*, which governs development of various neuronal subtypes^38^, were all significantly upregulated in WT organoids compared to eGBOs. In contrast, we observed genes implicated in GBM progression significantly upregulated in eGBOs compared to WT organoids (Fig. 2e). In particular, *APOE*^39^, *TMEM88*^40^, and *CXCL11*^41^ were significantly upregulated in PRO eGBOs compared to WT organoids. the significantly upregulated genes in MES eGBOs compared to WT organoids included *APOE*^39^, *IFITM3*^42^, and *MDK*^43,44^. We also performed gene set enrichment analysis (GSEA) on differentially expressed genes with a Log_2_(Fold Change) > 1.0. This analysis revealed that genes upregulated in WT organoids were significantly enriched in gene sets across multiple lists from the Molecular Signatures Database (MSigDB) associated with normal neural development, while genes upregulated in eGBOs were significantly enriched in gene sets associated with cancer pathways (Fig. 2f).

To better understand the underlying networks driving the observed transcriptional differences between WT organoids and eGBOs, we performed inferred gene regulatory networks (iGRN) analysis (Fig. 2g, Extended Data Fig. 3). This method quantifies transcription factor (TF) activity by constructing regulons, which consist of a transcription factor and its downstream target genes based on co-expression inferencing^45,46^. We identified 411 regulons and observed distinctive patterns in regulon activity between WT organoids and eGBOs (Supplementary Table 3). UMAP clustering based on iGRN activity further delineated the transcriptional differences between WT organoids and eGBOs (Extended Data Fig. 3a). Notably, highly active regulons specific to WT organoids include TFs associated with brain patterning in developing organoids^47,48^, including SOX2(+), POU3F3(+), and PAX6(+) (Fig. 2g, Extended Data Fig. 3b), while TFs shown to be synergistic master regulators that promote tumorigenesis and invasion in gliomas^49^, such as STAT3(+) and C/EBPβ(+), were among the highly active regulons specific to both the PRO and MES eGBOs (Fig. 2g, Extended Data Fig. 3b). Interestingly, the most variable regulon within this dataset was PITX2(+), which is required for normal neuron development^50^ and associated with poor survival in GBM patients^51^. Further, high regulon activity for TFs that have been implicated in cancer stem cells was observed in both eGBOs. MES eGBOs had high activity of the HMGA2(+) regulon, which promotes GBM stemness, invasion and in vivo tumor formation^52,53^ (Fig. 2g), while PRO eGBOs had high activity of the FOXJ3(+) regulon, which is a neuroectodermal TF recently identified as a GBM-driving gene^54^. Together, these results demonstrate that underlying differences in gene regulatory networks alter the neural development of eGBOs and promote an oncogenic transcriptional landscape.

### Cellular architecture is disrupted in developing eGBOs

To understand how the altered transcriptional landscape of eGBOs affects spatial organization in brain organoids, we performed in situ sequencing (ISS) using the Xenium platform (10x Genomics) to generate high-resolution spatial gene expression maps using a targeted panel of 266 genes (Supplementary Table 4). This enabled spatial localization of genes critical for neural development^55-58^ and GBM progression^59-61^, such as *NES, PAX6, MEIS2, SOX11*, *SOX2*, *SOX9*, *EGFR* and *CDH1* (Extended Data Fig. 4a).

While ISS provides high spatial resolution of gene expression, the limited panel of target genes creates challenges for robust downstream analyses, such as cell type annotation. Therefore, we applied the anchor-based integration workflow, implemented in Seurat^62^, to perform probabilistic transfer of cell type annotations from scRNAseq data generated from batch-matched organoids. Visualization of cell type annotations in WT organoids revealed an expected morphological arrangement, with radial glia cells located toward the center of the developing organoid, surrounded by proliferating and neuronal-like cells (Fig. 3a). In both PRO and MES eGBOs this architecture is disrupted, with less clear delineation of developing neuronal regions around radial glial populations (Fig. 3a). Interestingly, the proportions of radial glial, mesenchyme, and neuronal cells were similar in WT organoids and eGBOs (Fig. 3b). However, a greater proportion of off-target cell types was observed in eGBOs, especially the MES eGBO, which had much larger neuroepithelial, neural crest, retinal, and microglia populations.

**Fig. 3:**
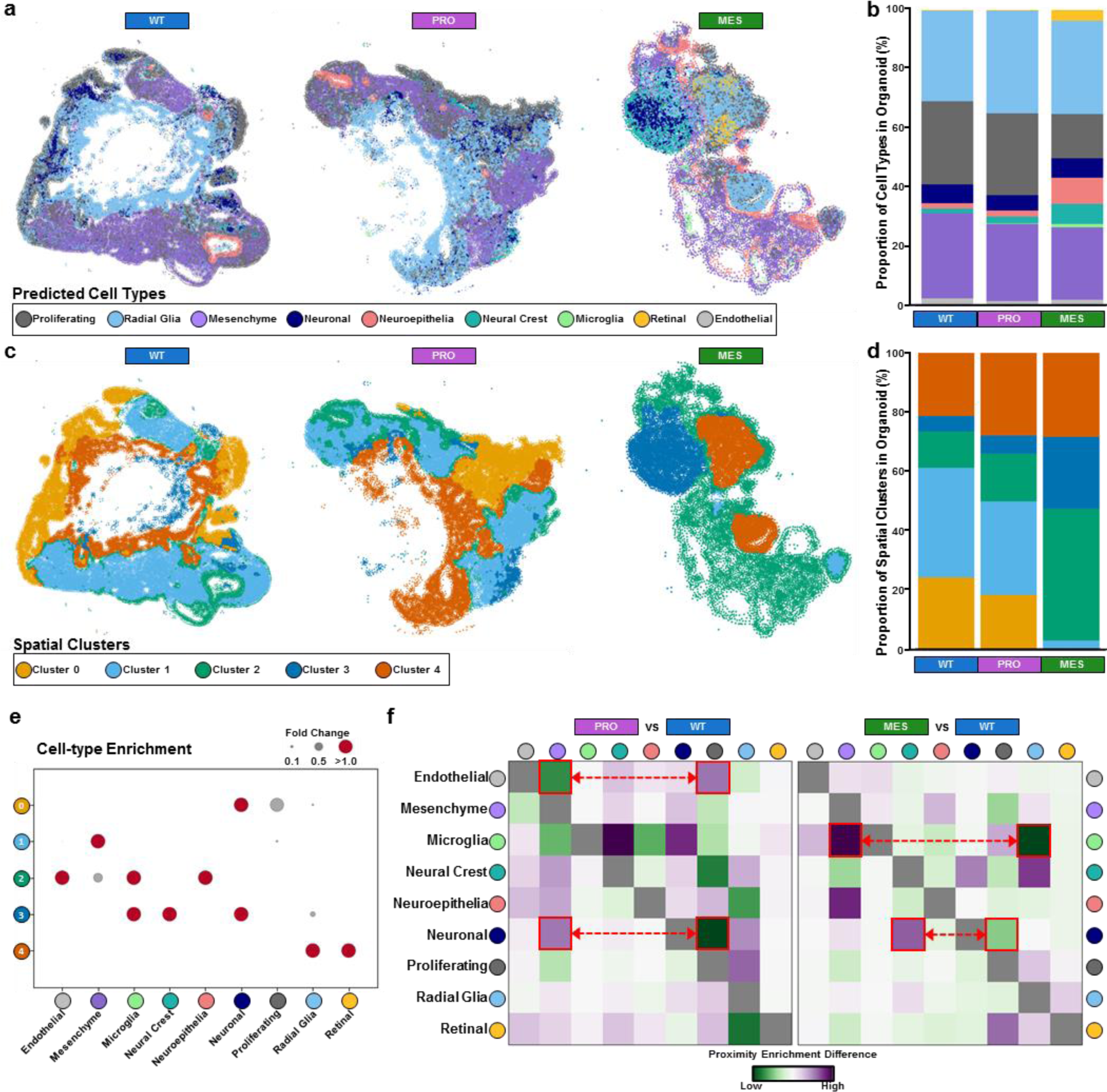
Spatial sequencing analysis of eGBOs. **a,** Spatial distribution of cell types predicted from scRNAseq data in WT, PRO, and MES organoids. **b,** Stacked bar plot showing the proportion of cell types in WT, PRO, and MES organoids. **c,** Spatial distribution of clusters determined by cluster stability analysis. **d,** Stacked bar plot showing the proportion of spatial clusters in WT, PRO, and MES organoids. **e,** Cell-type enrichment in each spatial cluster. **f,** Differential neighborhood enrichment heatmap between spatial distribution of major cell types in WT and eGBOs. Examples of differentially enriched neighborhoods are highlighted in red.

Next, we performed spatial clustering to define regions based on the transcriptional profile of both individual cells and cells in the surrounding neighborhood^63^. We identified five spatially defined clusters, based on cluster stability analysis, across both WT organoids and eGBOs (Fig. 3c, Extended Data Fig. 4b). Interestingly, WT organoids and PRO eGBOs consisted of similar proportions of each spatial cluster (Fig. 3c-d). In contrast, MES eGBOs consisted of larger proportions of Cluster 2 and Cluster 3 and decreased proportions of Cluster 0 and Cluster 1 (Fig. 3c-d). To characterize the composition of spatially defined clusters, we calculated cell-type enrichments within each spatial cluster. Expectedly, each spatial cluster was associated with a unique combination of cell types (Fig. 3e). Cluster 0 clearly marked a developing neuronal region, indicated by enrichment of neuronal and proliferating cell types. Interestingly, this cellular niche was absent from MES eGBOs and slightly reduced in PRO eGBOs. Cluster 1, marking primarily mesenchymal cells, was also much smaller in MES eGBOs and slightly reduced in PRO eGBOs. Multiple cell types, mostly off-target cell populations, were enriched in Cluster 2 and Cluster 3 and comprised a larger proportion of the eGBOs than they did in WT organoids. Additionally, Cluster 4, which primarily consisted of radial glial cells, was slightly larger in both eGBOs compared to WT organoids.

To determine changes in the spatial proximity of the different cell populations between WT organoids and eGBOs, we performed neighborhood enrichment analysis (Extended Data Fig. 4c). We also, compared the differences in neighborhood enrichment between eGBOs and WT organoids across cell populations (Fig. 3f). In PRO eGBOs, neuronal cells decreased interaction with proliferating cells, in favor of interactions with mesenchymal cell types, compared to WT organoids. Additionally, there was an increased interaction of endothelial cells with proliferating cells, suggesting pro-angiogenic signaling, compared to WT organoids. In MES eGBOs, neuronal cells decreased interaction with proliferating cells and instead interacted more closely with neural crest cells, compared to WT organoids. Also, microglia cells strongly shift away from radial glial, favoring interaction with mesenchymal cells, in MES eGBOs compared to WT organoids. Together, these data demonstrate that the cellular architecture of developing brain organoids is altered in eGBOs.

### eGBOs form glioma-like tumors in vivo

To assess if the observed transcriptional and architectural differences of eGBOs were indicative of tumor initiating cells, we tested the oncogenic potential of these cells in vivo. Approximately 2.6×10^5^ cells were stereotactically injected into the base of the right forebrain of athymic mice (Fig. 4a). Magnetic resonance imaging (MRI) was performed to track tumor formation and growth. Clear contrast-enhancing lesions were observed in mice that were injected with eGBO cells at three months post-transplantation (Fig. 4b). Hematoxylin & Eosin (H&E) staining in harvested brains showed extensive glioma-like growths in both PRO and MES eGBO conditions (Fig. 4c). We utilized a deep learning model (https://gbm360.stanford.edu/)^64^ to characterize cellular heterogeneity and predict prognosis based on the histology of observed growths (Extended Data Fig. 5a). In both PRO and MES eGBO conditions, the machine learning algorithm identified regions of tumor cells corresponding to the observed glioma-like growths. The PRO tumor consisted primarily of neural-progenitor-like (NPC-like) GBM cells, while the MES tumor consisted of a mix of NPC-like, oligodendrocyte-progenitor-like (OPC-like), astrocyte-like (AC-like), and mesenchymal-like (MES-like) cells. Further, the model assigned a moderate to high aggressiveness score to the cells in these regions.

**Fig. 4:**
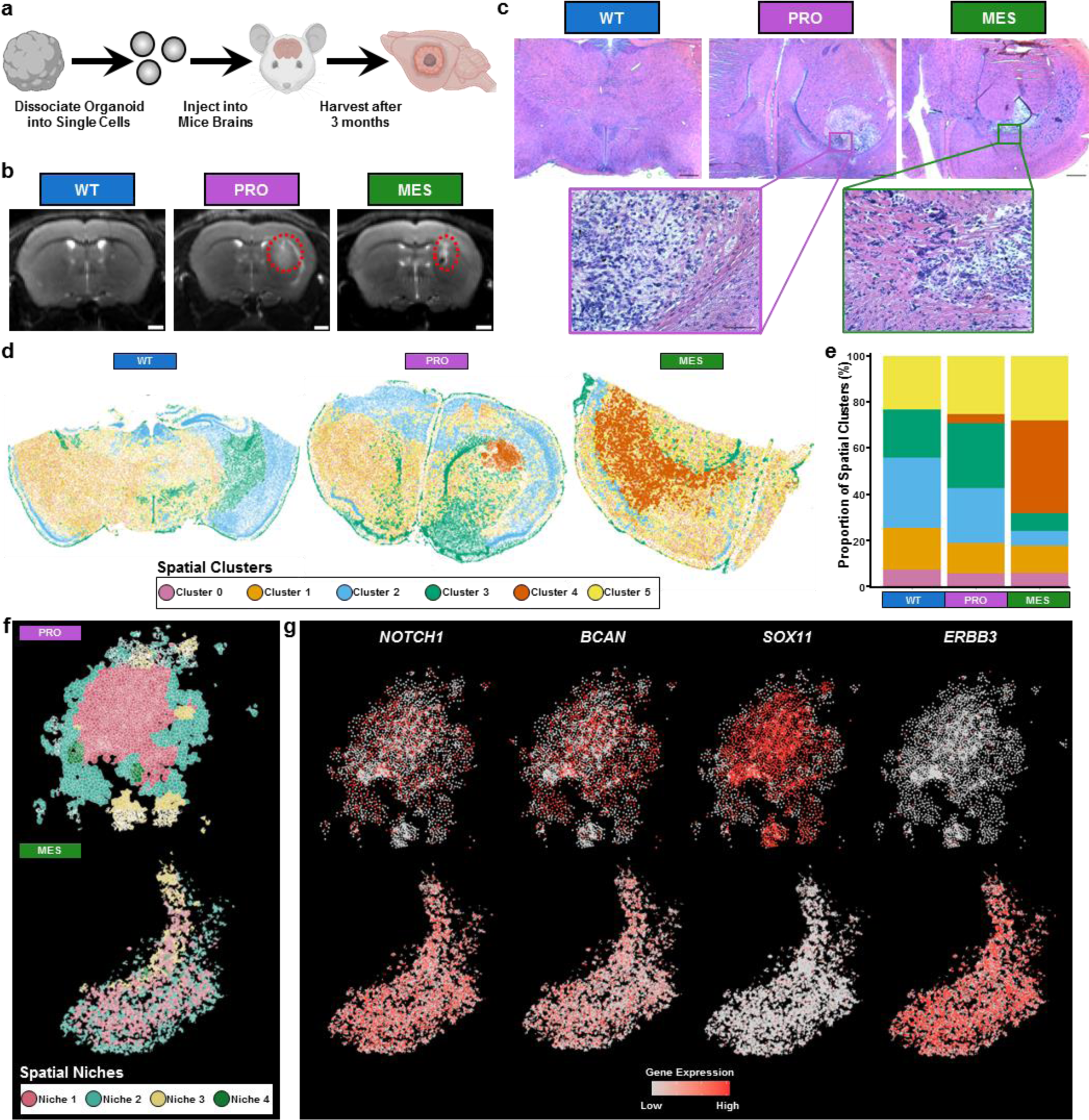
Spatially resolved transcriptomics analysis of eGBO-derived tumors. **a,** Experimental plan to assess tumorigenicity of eGBOs. Created using BioRender.com icons. **b,** Representative T2-post contrast MRIs of mouse brains 3 months post-transplantation of dissociated eGBO cells. Red, dotted circles indicate contrast-enhancing lesions; Scale bars = 1mm **c,** H&E staining of brain sections demonstrating glioma-like growths in mice that received cells from PRO and MES eGBOs (Scale bars = 500µm). Insets highlight characteristic histopathology of high-grade gliomas (Scale bars = 100µm). **d,** Spatial distribution of clusters determined by cluster stability analysis in recovered mouse brains. **e,** Stacked bar plot showing the proportion of spatial clusters identified in recovered mouse brains. **f,** Spatially defined niches within tumor regions identified in mice that received cells from PRO and MES eGBOs. **g,** Spatial distribution of genes associated with GBM development and migration in eGBO-derived tumors.

We performed ISS on recovered mouse brains using the 10x Xenium Human Brain panel to characterize the spatial transcriptional profile of observed growths. There are thirty-four tumor-associated genes in the 10x Xenium Human Brain panel. The localization of these transcripts identified tumor regions in both eGBO conditions (Extended Data Fig. 5b). To further characterize spatial patterning of gene expression, we calculated the Moran’s I global spatial auto-correlation statistics for all genes (Supplementary Table 5). Comparing genes with a Moran’s I statistic greater than 0.1 revealed that eight genes were spatially autocorrelated in both eGBO conditions (Extended Data Fig. 5c). Among these shared genes were the most highly spatially autocorrelated genes, *NNAT* and *IGFBP3*, in the PRO and MES conditions, respectively, which have both been associated with GBM progression and reduced patient survival^65,66^.

Spatial clustering revealed six regions across brain sections recovered from mice injected with cells derived from WT, PRO, or MES organoids (Fig. 4d, Extended Data Fig. 5d). These clusters represented various brain regions including cortical layers, indicated by expression of *CUX2*, *RORB*, and *LAMP5*, and the dorsal thalamus, marked by expression of *NTNG1*^67^ (Extended Data Fig. 5e). Notably, Cluster 4, which was present only in mice injected with cells from eGBOs, clearly marked regions aligning with glioma-like growths observed in MRI scans and H&E staining (Fig. 4d-e).

To further characterize the spatial transcriptomic profile of tumors generated by eGBOs, we computed spatial niches within the spatial cluster corresponding to glioma-like growths, revealing distinct spatial organization in PRO and MES eGBO-derived tumors (Fig. 4f). Recent integrative spatial analysis demonstrated that human GBM consists of both structured and disorganized regions^68^. Based on the distribution of spatial niches in eGBO-derived tumors, we find that the PRO eGBO-derived tumor has a more structured organization, while the MES eGBO-derived tumor was more disorganized. Both PRO and MES eGBO-derived tumors had broad expression of *NOTCH1*, which is commonly upregulated in glioma tumor-initiating cells^69^, and *BCAN*, which encodes an extracellular matrix protein found in human gliomas^70^ (Fig. 4g). However, expression of *SOX11*, a TF crucial for brain development that is also overexpressed in GBM^71^, appeared to be localized to the core of the PRO eGBO-derived tumor, while the MES eGBO-derived tumor broadly expressed *ERBB3*, a subtype-specific GBM marker^72^. Together, these data demonstrate that eGBOs can generate tumors that demonstrate subtype-specific characteristics of human GBM tumors.

### eGBO-derived tumors recapitulate cell states found in human GBM

To characterize the transcriptional profile of the eGBO-derived tumors more thoroughly, we performed scRNAseq on dissociated cells isolated using a brain tumor dissociation kit. Substantially fewer single cells were recovered from mice that received cells from WT organoids compared to mice that received cells from eGBOs. Additionally, the mean genes per cell from cells recovered from mice treated with WT organoid cells was greatly reduced compared to cells recovered from mice treated with eGBOs. These data suggest mouse cells were most likely recovered from the WT organoid condition, due to the low mapping of human genes, and further suggest that WT organoids do not form glioma-like growths in vivo. Following quality control filtering, 476 high-quality cells were recovered from transplanted WT organoids while 10,103 high-quality cells and 12,452 high-quality cells were recovered from transplanted PRO and MES eGBOs, respectively, for further analysis.

Cancer cells are difficult to classify as distinct cell types^73-76^, therefore we integrated our scRNAseq data with previously published data from twenty-nine patient GBM samples^77^ to characterize eGBO-derived tumor cells (Fig. 5a, Extended Data Fig. 6a). Non-malignant cells were identified based on previously reported marker genes^77^ corresponding to immune cells and oligodendrocytes (Fig. 5b, Supplementary Table 6). Quantification of the proportion of each sample across malignant and non-malignant cell types revealed that the non-malignant cell types consisted primarily of non-malignant patient cells and cells recovered from transplanted WT organoids while the majority cells recovered from transplanted eGBOs were classified as malignant (Fig. 5c). Pearson correlation analysis also demonstrated that the transcriptional profiles of eGBO-derived tumor cells were more similar to malignant patient cells, while recovered WT organoid cells were more similar to non-malignant patient cells (Fig. 5d).

**Fig. 5:**
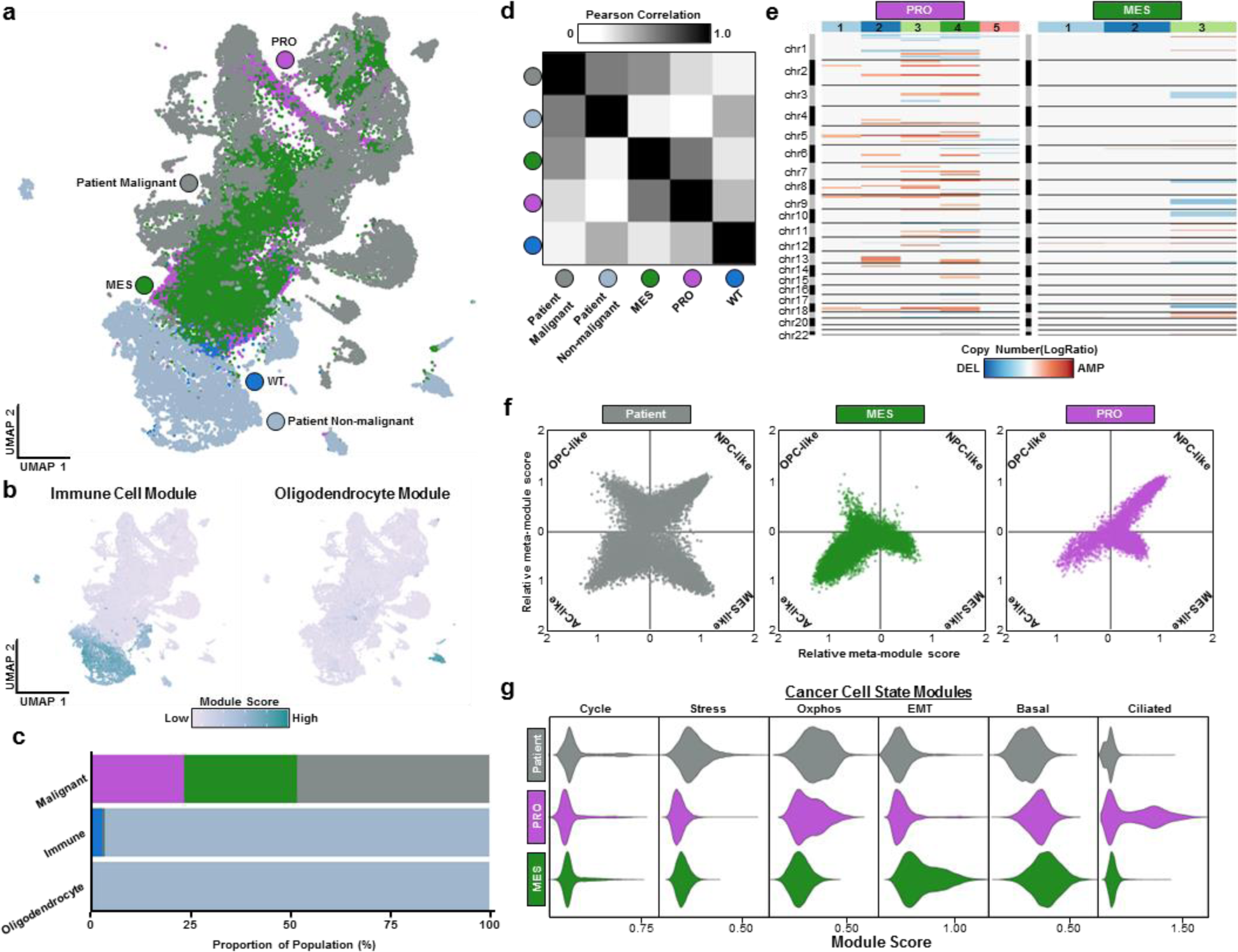
Integrative transcriptional profiling of eGBO-derived tumors and patient GBM samples. **a,** UMAP of integrated recovered WT organoid cells, eGBO-derived tumors, and GBM patient samples. **b,** Feature plots indicating module scores for non-malignant cell types present in GBM samples. **c,** Stacked bar plot showing the distribution of each sample across malignant and non-malignant cell types. **d,** Compact representation of CNVs present in each subclone identified in eGBO-derived tumors. **e,** Heatmap of Pearson correlation coefficient for 2000 most variable expressed for WT organoid cells, eGBO-derived tumors, and GBM patient samples. **f,** Two-dimensional representation of malignant cell states. Each quadrant corresponds to a cellular state and the position of each dot reflects the relative meta-modules scores for individual malignant cells. OPC-like = oligodendrocyte-progenitor-like; NPC-like = neural-progenitor-like; MES-like = mesenchymal-like; AC-like = astrocyte-like. **g**, Violin plots indicating module scores for recurrent cancer cell states in malignant cells from eGBO- and patient-derived tumors.

Copy-number variations (CNV) are a hallmark of malignant glioma cells, with sixteen broad events identified in GBM samples^78^. We assessed the landscape of CNVs in eGBO-derived tumors and patient-derived GBM samples using a variational algorithm to characterize the copy number profile of clonal substructures within tumors from scRNAseq data^79^ (Fig. 5d, Extended Data Fig. 6b). Notably, we observed subclones within the PRO eGBO-derived tumor cells with amplification in chromosome 7, while a deletion on chromosome 10 was observed in one of the MES eGBO-derived tumor cell subclones. These observations are consistent with prior assessments in patient samples^77,78^, that found alterations in chromosomes 7 and 10 were most common despite high inter-tumoral variability between patients and intra-tumoral variability within subclones.

Prior characterization of inter-tumoral and intra-tumoral heterogeneity in patient GBM samples identified four recurrent cell states that occur across GBM cells^77^. Consistent with these prior findings, we observe not only a heterogenous distribution of cell states across patient-derived tumor cells, but also mutation-specific distributions of cell states in eGBO-derived tumor cells (Fig. 5f). MES eGBO-derived tumor cells existed predominantly in the AC-like state, with some cells falling in the OPC-like or MES-like states. PRO eGBO-derived tumor cells were largely in the NPC-like state, with some cells associated with the MES-like or AC-like states.

In addition to these GBM-specific cell states, recurrent cell states that constitute general features of cancer have been identified^73^ (Supplementary Table 6). Analyzing the module score of these recurrent cell states, we found that eGBO-derived tumors recapitulate some aspects of patient-derived GBM tumors (Fig. 5g), including a subset of cells expressing cell cycle genes (Cycle module), stress genes (Stress module), and oxidative phosphorylation genes (Oxphos module). Interestingly, we also observe subtype-specific enrichment of modules, such as the epithelial-mesenchymal transition module (EMT) in MES eGBO-derived tumor cells and the ciliated epithelial cell module (Ciliated) in PRO eGBO-derived tumor cells. These data indicate tumors formed by eGBOs recapitulate the heterogenous composition of cell states found in human GBM.

### Developmental states in eGBOs model cancer cell states found during GBM progression

Recently, it has been proposed that cancer cell states in GBM and other cancers are constrained within the developmental hierarchy of the cell that originates tumor formation^16,18,80,81^. Our eGBOs combined with patient tumor samples offer a unique model to investigate this hypothesis. Therefore, we generated two integrated datasets consisting of eGBOs, eGBO-derived tumors, and malignant patient-derived GBM samples with comparable mutational profiles (Fig. 6a, Extended Data Fig. 6c).

**Fig. 6:**
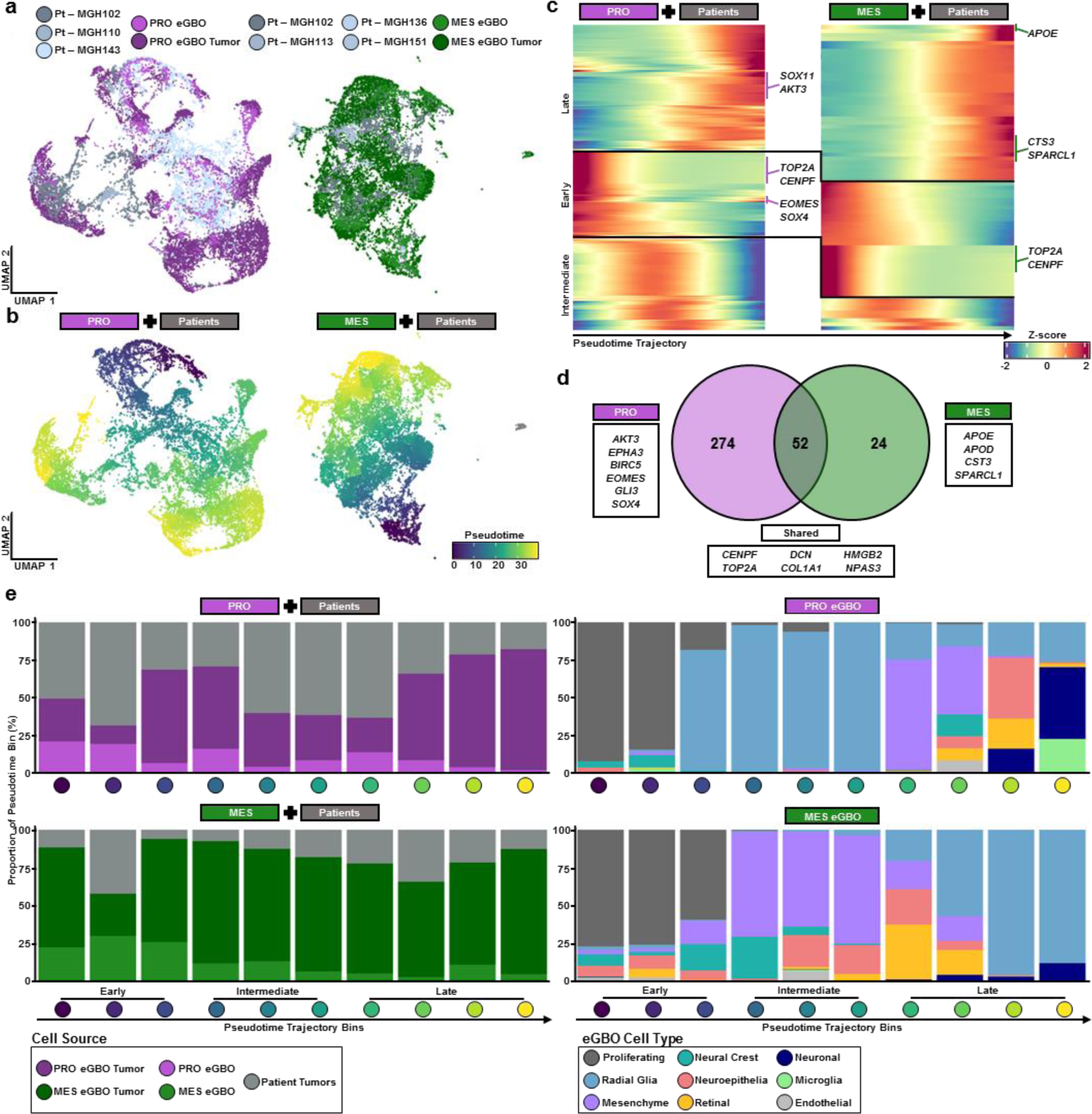
Trajectory analysis reveals developmental states govern GBM progression. **a,** UMAPs of integrated datasets for eGBOs, eGBO-derived tumors, and GBM patient samples. The PRO dataset consists of PRO eGBOs, PRO eGBO-derived tumor cells, and cells derived from GBM patients with a *TP53* mutation. The MES dataset consists of MES eGBOs, MES eGBO-derived tumor cells, and cells derived from GBM patients with a *PTEN* mutation. **b,** UMAPs showing the pseudotime trajectory of cells from a proliferating population to different cancer cell states for PRO and MES integrated datasets. **c,** Trajectory heat maps showing changes of gene expression across pseudotime for PRO and MES integrated datasets. **d,** Venn diagram of pseudotemporally autocorrelated genes from PRO and MES integrated datasets. **e,** Stacked bar plots showing the proportion of cells from eGBOs, eGBO-derived tumors, and GBM patient samples (left) and the proportion of eGBO cell types (right) across pseudotime bins.

We performed single-cell trajectory analysis to characterize the changes in gene expression as proliferative cells transition to malignant cell states (Fig. 6b). Clustering of genes that have expression correlated to pseudotime trajectory revealed three distinct transcriptional stages (Fig. 6c,). Interestingly, we found a larger number of genes correlated with pseudotemporal stages in the PRO integrated dataset compared to the MES integrated dataset (Fig. 6d). Shared pseudotemporally correlated genes between both integrated datasets included genes associated with proliferation (*CENPF* and *TOP2A*), cancer-associated fibroblasts^82,83^ (*DCN* and *COL1A1*) and neurodevelopment^84,85^ (*HMGB2* and *NPAS3*). Pseudotemporally correlated genes specific to MES integrated dataset included differentiation markers found in astrocytes or oligo-lineage cells, such as *APOE* and *APOD*, that also mediate pro-inflammation and proliferation signatures in GBM^18,86^. Genes associated with GBM tumor-initiating cells^87-89^ (*AKT3, EPHA3, BIRC5,* and *EOMES*) and radial glia cells (*GLI3* and *SOX4*) were unique to the pseudotemporally correlated gene list of the PRO integrated dataset. To investigate changes in cell states across tumor progression, we binned cells based on their pseudotime ordering and quantified the proportion of cell types in each bin (Fig. 6e). The distribution of cell types across the pseudotime further delineated similarities and differences between the PRO and MES integrated datasets. In both datasets, eGBO cells along with patient- and eGBO-derived tumor cells were well distributed across the pseudotime ordering. Additionally, the early pseudotime stages in both datasets consisted of primarily of proliferating cells from eGBOs prior to transplantation. Notably, the cell populations in the intermediate and late pseudotime stages appear to be swapped in between the two datasets. Specifically, radial glial cells from eGBOs appear in the intermediate pseudotime stage in the PRO dataset, while mesenchymal and other cell types are present in the late pseudotime stage. In the MES dataset, on the other hand, mesenchymal and other cell types appear in the intermediate pseudotime stage, while radial glial cells from eGBOs are primarily found in the late pseudotime stage. This is consistent with previous observations in human GBM samples demonstrating that proliferating GBM stem cells exist on an axis of PRO-MES, which contributes to the heterogeneity observed in GBM^90^. Additionally, the presence of cells from eGBOs and both patient- and eGBO-derived tumors in the late pseudotime stage of suggests that early developmental states observed in eGBOs are representative of malignant cell states that arise during GBM progression. Together, these data highlight that subtype-specific eGBOs can model different developmental states that contribute to the heterogeneity of GBM during tumor progression.

## DISCUSSION

The lack of effective treatments for GBM highlights the need for improved preclinical models that replicate human tumorigenesis and progression. By introducing GBM subtype-specific mutations using genetic engineering methods, we have developed eGBOs as an in vitro model system for GBM. The combination of brain organoids and genetically engineered oncogenic aberrations enabled us to characterize the disruption of developmental pathways and the progression of cancer cell states that are characteristic not only of GBM, but also a broad range of cancers. Further, GBM subtype-specific mutations within eGBOs enabled characterization of the progression of different GBM subtypes through developmental states.

Prior genetically engineered organoid models of brain tumors have exhibited many features of cancer, including malignant cellular identities and the capacity for in vivo expansion and invasion^8,19,20^. Induced expression of the oncogenic HRas^G12V^ mutation, which is not common in patients, was demonstrated as a proof of principle that cerebral organoids could be used as a model for tumor formation^19^. Similarly, inducing specific combinations of oncogene expression or mutations via nucleofection promoted GBM-like overgrowths within developing organoids that could be utilized for drug screening^20^. Building on this work, we targeted the *TERT* promoter region to create a defined genetic background that recapitulates one of the earliest GBM mutations that provides a selective advantage for malignant cells^10^. We further generated subtype-specific eGBOs by targeting some of the common mutations found in subsets of patient samples. Orthotopic transplantation of eGBO cells resulted in tumors that exhibited many characteristics of GBM, including histological presentation, spatial organization, and cell state heterogeneity. We observed distinct differences in spatial architecture and cell states in tumors formed by PRO and MES eGBOs that can be attributed to the differences in their respective genetic backgrounds. Notably, these differences correspond to characteristics of human GBM samples that contribute to inter- and intra-tumor heterogeneity.

While eGBOs are a promising model for further elucidating human brain tumor biology and screening tool for potential therapeutic approaches, they still lack components that are critical for tumor progression. For example, existing vasculature is involved in multiple GBM processes, such as glomeruloid body formation^91^ and pseudopalisading necrosis^92^. Interestingly, a small population of endothelial cells was present in eGBOs suggesting that they may be suitable model to study angiogenic signaling during GBM progression. integration of eGBOs and vascularized organ-on-chip models can also enable investigation of early tumor interactions with existing vascular networks and tumor invasion. Additionally, many genes are mutated in GBM, and the mutational burden can vary significantly not only between patients, but also within patient samples. While we only explored a small subset of GBM-associated mutations, our results demonstrate that eGBOs can be utilized to interrogate how specific mutation combinations drive different features of GBM.

Our single-cell and spatial transcriptomic analyses of eGBOs provides an important validation of engineered cancer organoid models and demonstrate their utility as a human-relevant model of GBM tumorigenesis for future preclinical drug development. Specifically, we show that eGBOs can model different aspects of GBM progression based on specific combinations of mutations and future development can focus on investigating the pathogenesis of different, patient-specific mutations. This model offers a complementary approach to existing methods for investigating tumor biology and pathophysiology. Ultimately, eGBOs have the potential to enable early drug discovery of therapeutics designed to target patient-specific genetic backgrounds.

## METHODS

### Ethics Statement

Human stem cell research related to this study has been approved by the Washington University School of Medicine Institutional Review Board and Embryonic Stem Cell Research Oversight Committee (IRB ID: 201709124 and ESCRO# 17-005). The Washington University School of Medicine Institutional Animal Care and Use Committee (IACUC) approved the protocols for experiments involving animals.

### Culture and Genetic Engineering of hPSCs

H1 hPSCs (CVCL_9771) were maintained on Matrigel-coated T75 culture flasks in mTeSR1 media (Stemcell Technologies; 85850). During passaging, media was supplemented with 10µM Y-27632 (Abcam; ab120129) to inhibit Rho kinase-mediated apoptosis. Genetically engineered H1 hPSCs cell lines were generated by the Genome Engineering & Stem Cell Center at Washington University School of Medicine in St. Louis. Briefly, H1 hPSCs were transfected with Cas9 recombinant protein and synthetic single guide RNA to introduce a C>T mutation within the telomerase reverse transcriptase (*TERT*) gene promoter^21^. The H1 TERT mutant cell line was then further to modified, by the same procedure, to introduce either a point mutation targeting the *TP53* gene or knockouts mutations in the *PTEN* and *NF1* genes. CRISPR guide RNA sequences are listed in Supplementary Table 7.

### Immunocytochemistry

For immunocytochemistry (ICC), cells were fixed in 4% paraformaldehyde (Electron Microscopy Science; 15714) for 30min at room temperature (RT). Fixed cells were then incubated at RT in ICC solution consisting of 0.1% Triton X (Acros Organics; 327371000) and 5% donkey serum (Jackson Immunoresearch; 01700-121) in phosphate buffered saline (Fisher; MT21040CV), for 30min to permeabilize cell membranes and block non-specific binding sites. Samples were subsequently treated with primary (R&D Systems; AF1997) and secondary (Invitrogen; A11055) antibodies in ICC solution overnight at 4°C and 2h at RT, respectively. DAPI (Invitrogen; D1306) was used for nuclear staining. Samples were incubated in DAPI for approximately 12min at RT, washed with ICC solution and stored in PBS until imaging.

### Quantitative PCR Analysis

RNA was extracted from cells using the RNeasy Mini Kit (Qiagen; Cat #74104) following the manufacturer’s instructions. Quantification and quality assessment of RNA was performed using a BioTek Synergy H1multimode microplate reader. Complementary DNA (cDNA) was synthesized from extracted RNA using the using the High-Capacity cDNA Reverse Transcription Kit (Applied Biosystems; 129382310MG) and a T100 thermocycler (Bio-Rad). The TaqMan™ Universal Master Mix (Applied Biosystems; 4440040) and predesigned TaqMan Gene Expression Assays (Applied Biosystems; 4331182) were used to detect mRNA transcript levels on a QuantStudio™ 6 Pro Real-Time PCR System (Applied Biosystems; A43180). Results were analyzed using the ΔΔCt methodology with GAPDH used as a housekeeping gene. Primers for qPCR are listed in Supplementary Table 8.

### Brain Organoid Differentiation

Brain organoids were generated from H1 hPSC lines following a previously described protocol^32^, with minor modifications. Briefly, H1 hPSCs were cultured in hPSC Media on Matrigel-coated T75 flasks. On day 0 of differentiation, cells were transferred into low-attachment 96-well plates at approximately 10,000 cells/well to generate embryoid bodies. On day 2, each embryoid body was transferred to a single well of a 24-well plate with Neural Induction Media. On day 6, developing neural organoids were embedded in Matrigel droplets and transferred to a petri dish with Cerebral Organoid Differentiation Media. On day 10, petri dishes were transferred to an orbital shaker at 100rpm and fed every four days with Organoid Differentiation Media. Media formulations and additional details are provided in Supplementary Table 1.

### Organoid Morphology Analysis

The Fiji distribution of ImageJ was used to perform area and circularity measurements^93^. Briefly, bright-field images were converted to 8-bit greyscale, then binarized to delineate the borders of the developing organoid. Edges of the organoid area was defined using the wand tracing tool. Area and shape descriptors were measured using the default Measure functions. Circularity was calculated as 4π(area/perimeter^2^), with a value of 1.0 indicates a perfect circle.

### Orthotopic Xenotransplantation

Dissociated brain organoid cells were injected into the brain of seven-week-old female athymic nude mice as previously described^19^. Briefly, approximately 260,000 cells were injected stereotactically into seven-week-old female athymic nude mice (n = 5 per condition). Injections targeted the right putamen at coordinates 1mm rostral to bregma, 2mm lateral, and 2.5mm deep. To track tumor growth, MRI was performed on a Bruker BioSpec 9.4T MRI (Bruker, Billerica, MA) using an 86mm inner diameter volume transmitter coil and a 4-channel mouse brain CryoProbe array receiver coil. Prior to imaging, mice were anesthetized with 1-1.5% isoflurane on a circulating warm water pad to maintain body temperature. Fat-suppressed T2-weighted RARE images were acquired of the brain using the following parameters: TR = 2500s, TE = 33s, matrix size = 256×256, spatial resolution = 700μm, 9 slices at 5mm thickness, RARE factor = 9, 1 average, acquisition time = 1min. Image analysis was done using the semi-automatic segmentation analysis software ClinicalVolumes (ClinicalVolumes, London, UK).

### Tumor Dissociation and Histology

Three months after transplantation, mice were sacrificed, and their brains were harvested. For scRNAseq analysis, tumors were dissociated using the Brain Tumor Dissociation Kit (Miltenyi Biotec, 130-095-942). Myelin was removed using Myelin Removal Beads II (Miltenyi Biotec, 130-096-731). Mouse cells were removed using the Mouse Cell Depletion Kit (Miltenyi Biotec, 130-104-694). Dissociated cells from two mice per condition were pooled together. The other three brains were processed for histology. Briefly, samples were fixed with 4% PFA cryopreserved with 30% sucrose, and flash frozen. Fixed samples were sectioned at 10μm thickness along the coronal plane using the tissue near the needle tract for cell injection based on stereotactic coordinates. Brain sections were stained with hematoxylin to mark the cell nuclei and eosin to stain the extracellular matrix and cytoplasm.

### Single-Cell RNA Sequencing

Cell suspensions were delivered to the Washington University in St. Louis Genome Technology Access Center (GTAC) for library preparation and sequencing. For library preparations, the Chromium Next GEM Single Cell 3ʹ Reagent Kit v3.1 (10x Genomics) was used following the provided protocol. The library was sequenced with a NovaSeq 6000 System (Illumina) at the recommended 26x98bp. To label cells with unique hashtags, approximately ten organoids per condition were pooled and dissociated in TrypLE for 15min with gentle agitation every 5min. Single cell solutions were resuspended in 200µL of cell staining buffer (BioLegend, 420201) plus 1µg of TotalSeq-A hashtag antibodies (BioLegend, 399907) and incubated for 30min on ice. After three washes in 1mL of cell staining buffer, cells were resuspended in DMEM at a concentration of 1000cells/µL and pooled together.

### In Situ Sequencing

ISS was performed at GTAC using Xenium platform (10x Genomics), following protocols provided by the company. Sections were prepared according to the “Xenium In Situ for FFPE-Tissue Preparation Guide” (CG000578 Rev C, 10X Genomics) protocol. Briefly, 5µm sections were cut and carefully placed onto the Sample Area of a Xenium slide (PN-1000465). The slides were then placed in a drying rack at room temperature for 30min to remove excess water, followed by a 3h incubation at 42°C on a Xenium Thermocycler Adapter plate positioned in a thermocycler (BioRad). Subsequently, the slides were stored overnight at room temperature with a desiccant for further drying. The following day, Xenium slides were processed following the “Xenium In Situ for FFPE-Deparaffinization and Decrosslinking” protocol (CG000580 Rev C, 10X Genomics). The Xenium slides were then assembled into Xenium cassettes (PN-1000566, 10X Genomics), which allow for the incubation of slides on the Xenium Thermocycler Adapter plate in a PCR machine with a closed lid for precise temperature control. The slides were processed using the “Xenium Slides and Sample Prep Reagents” kit (PN-1000460, 10X Genomics), beginning with an incubation in a decrosslinking and permeabilization solution at 80°C for 30min. The Xenium slides were then processed according to the “Xenium In Situ Gene Expression” user guide (CG000582 Rev D, 10X Genomics) for the remaining slide preparation steps. The slides were incubated at 50°C for 17h with the pre-designed gene expression probe set, “Xenium Human Brain Gene Expression”. This was followed by a series of wash and enzymatic steps, including a 30min post-hybridization wash at 37°C, a 2h ligation at 37°C, and a 2h amplification step at 30°C. After additional washes, the slides were treated with an autofluorescence quencher and a stained with DAPI.

Processed Xenium slides, assembled in Xenium cassettes, were subsequently imaged using the Xenium Analyzer, following the guidelines provided in the “Xenium Analyzer User Guide: (CG000584 Rev B, 10X Genomics).” The two Xenium slides/cassettes were then loaded into the instrument, initiating the Analyzer’s ‘sample scan’ process. This scan produced images of the fluorescent nuclei in each section. Upon completion, these images were utilized to determine the regions within the scan area to be included in the instrument’s comprehensive scan of the gene expression probe set.

### Single-Cell RNA and In Situ Sequencing Data Analyses

Datasets were analyzed using a custom containerized Linux environment consisting of JupyterLab v4.0.11, R v4.4.1, and Python v3.10.13. Initial processing and quality control filtering of scRNAseq and ISS data was performed using Seurat v5^62^. Unsupervised annotation of eGBO cell types was performed using SingleR^94^ and VoxHunt^47^. GSEA was performed using fgsea^95^. Inferred GRN analysis was performed using SCENIC^45,46^. Additional analysis of ISS data was performed using Scanpy^96^, Squidpy^97^, CellCharter^63^, and scvi-tools^98^. Inferred CNV analysis was performed using SCEVAN^79^.

### Statistical Methods

Statistical analyses were performed using GraphPad Prism 8.0.1 software (GraphPad Software, San Diego, CA). Statistical analyses used included parametric unpaired t-test using the Holm-Sidak method and two-way ANOVA with matching factors corrected for multiple comparisons using Dunnett’s multiple comparison test and the Geisser-Greenhouse correction without sphericity. A confidence level of 95% was considered significant with designations of *P < 0.05, **P < 0.01, and ***P < 0.001 (two-tailed). Results without statistical significance denoted can be assumed to be of no significant difference. All values were expressed as mean ± standard deviation.

## DATA AVAILABILITY

All scRNAseq and spatial transcriptomics data generated in this study will be deposited in the Gene Expression Omnibus (GEO).

## CODE AVAILABILITY

Original codes to reproduce analysis and figures will be deposited on GitHub at https://github.com/ishahakm/eGBO. The custom docker image with all software packages for analyzing scRNAseq and spatial sequencing data is available at https://hub.docker.com/r/ishahakm/aussie.

## ACKNOWLEDGEMENTS

This work was funded by the Edward J. Mallinckrodt Foundation (to J.R.M.); startup funds from the Washington University School of Medicine Department of Medicine (to J.R.M.); the National Institutes of Health (NIH) R01 NS094670, R01 NS106612, R01 NS128470 (to A.H.K.); Hope Center for Neurological Disorders Pilot Grant (to A.H.K. and J.R.M); the Alvin J. Siteman Cancer Center Siteman Investment Program through funding from The Foundation for Barnes-Jewish Hospital (to A.H.K.); the Christopher Davidson and Knight Family Fund (to A.H.K.); the Duesenberg Research Fund (to A.H.K.). M.I. was supported by the Rita Levi-Montalcini Postdoctoral Fellowship in Regenerative Medicine and the NIH (T32DK007120). R.H.H. was supported by the NIH (R25 NS090978), American Cancer Society (PF-21-149-01-CDP), and Glioblastoma Foundation. We thank members of A.H.K. and J.R.M. labs for helpful discussions. We would also like to thank Erika Brown (Washington University) for helpful feedback on the manuscript. We would like to thank Drs. Yi-Hsien Chen, Yong Miao, Vijayalingam Selvamani, and Xiaoxia Cui (Washington University School of Medicine Genome Engineering and Stem Cell Center) who performed the CRISPR/Cas9 screening experiments for mutant hPSC generation. We would also like to thank Kymberli May (Washington University in St. Louis Advanced Imaging and Tissue Analysis Core of the Digestive Disease Research Core Center) who prepared the Xenium slides. The MRI studies presented in this work were conducted in the Small Animal MR Facility of the Mallinckrodt Institute of Radiology at Washington University supported by S10OD026913. We thank the GTAC staff at the McDonnell Genome Institute at Washington University School of Medicine for help with genomic analysis. The Center is partially supported by NCI Cancer Center Support Grant P30CA91842 to the Siteman Cancer Center from the National Center for Research Resources (NCRR), and NIH Roadmap for Medical Research. This publication is solely the responsibility of the authors and does not necessarily represent the official view of NCRR or NIH.

## AUTHOR CONTRIBUTIONS

M.I., A.H.K., and J.R.M. designed the project and experiments. M.I. performed in vitro experiments and performed the computational analyses. D.A., P.A.D., and X.Q. generated CRISPR-edited stem cell lines. R.H.H., C.M., R.T.C., and T.W. performed in vivo experiments. M.I. and S.D. performed assessment of histological samples. M.I. wrote the manuscript. All authors revised and approved of the manuscript.

## COMPETING INTERESTS

M.I. has stock in Vertex Pharmaceuticals. A.H.K. is a consultant for Monteris Medical and has received research grants from Stryker for a clinical outcomes study about a dural substitute, which have no direct relation to this study. J.R.M. was employee of and has stock in Sana Biotechnology. The remaining authors declare no competing interests.

## EXTENDED DATA FIGURES

**Extended Data Fig. 1:**
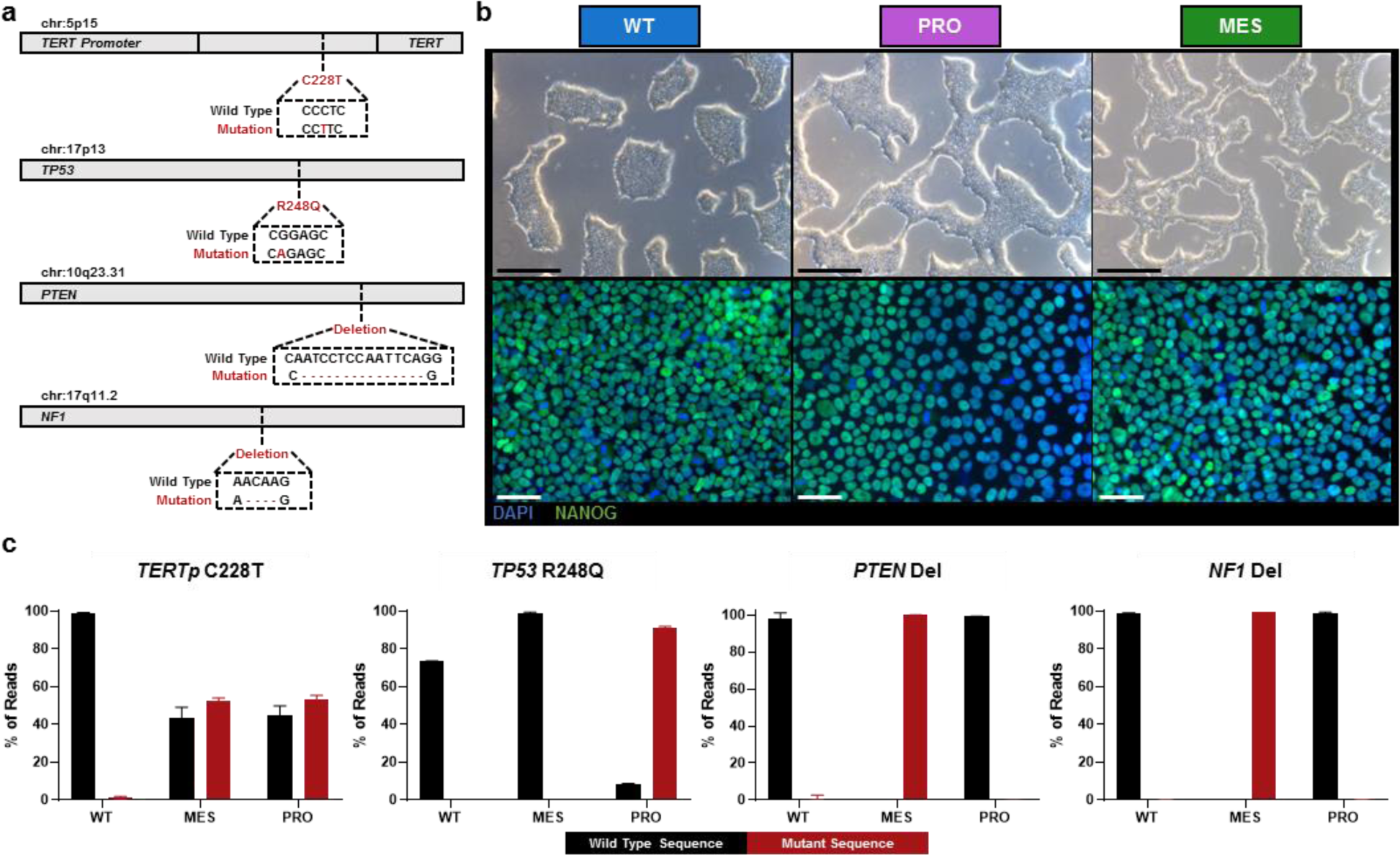
Characterization of stem cells harboring GBM-associated mutations. **a**, Depiction of genetic mutations introduced into hPSCs. **b**, Brightfield (top, scale bars = 500µm) and immunofluorescence imaging (bottom, scale bars = 50µm) of hPSCs indicating normal stem cell morphology and expression of the pluripotency marker, NANOG. **c**, NGS results demonstrating successful gene editing in engineering hPSCs (n = 4).

**Extended Data Fig. 2:**
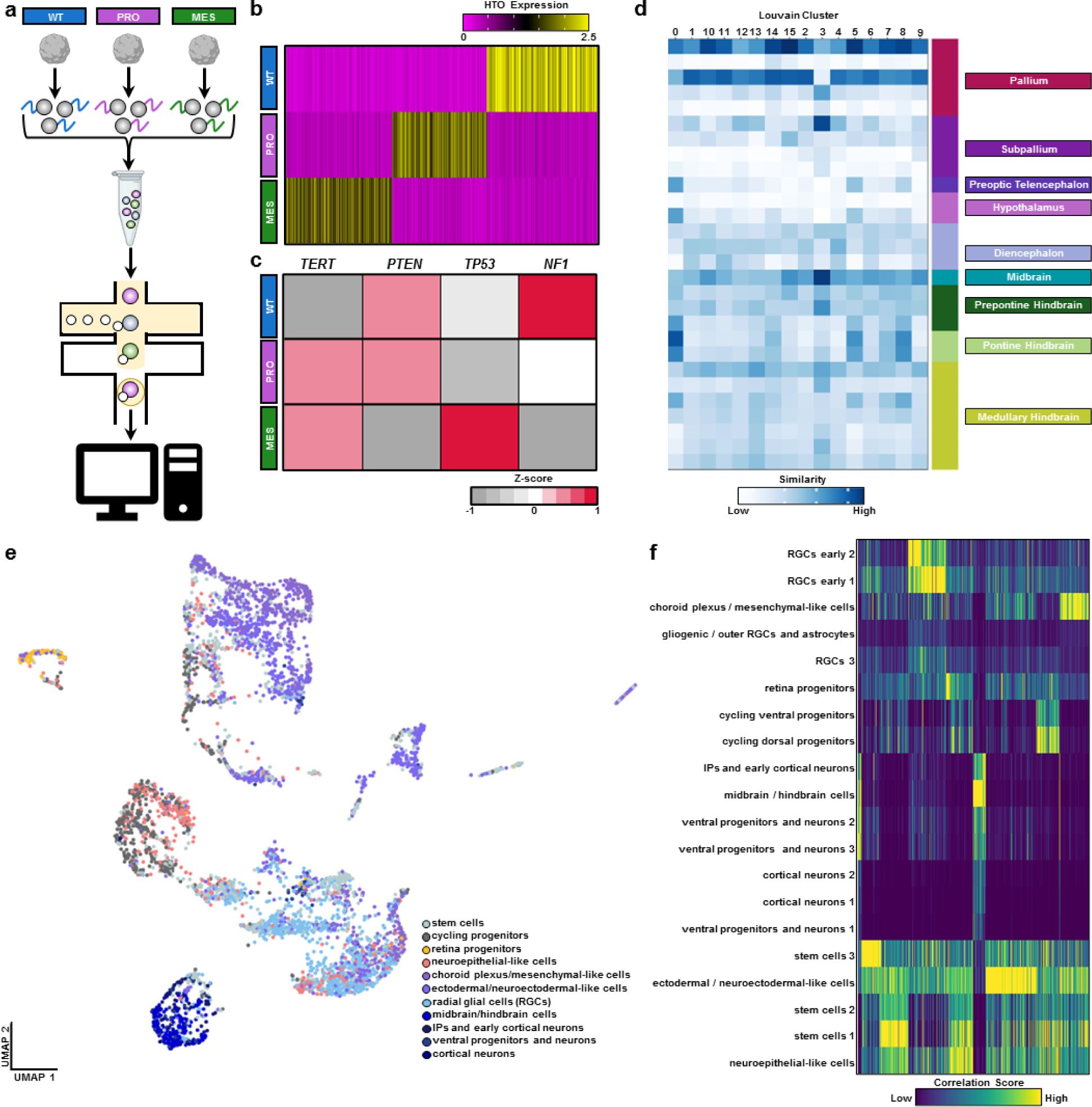
Hashtag demultiplexing and unsupervised cell type annotation in scRNAseq data. **a**, Schematic of multiplexed scRNAseq workflow. Briefly, organoids are dissociated into single cells and labeled with a unique hashtag oligonucleotides (HTO). Samples are then pooled and scRNAseq is performed. Finally, samples are demultiplex computationally. Created using BioRender.com icons. **b**, Heatmap showing expression of HTOs correctly defines each condition. **c**, Heatmap of genes targeted by CRISPR editing to mimic GBM subtypes. **d**, Heatmap of similarity index calculated by VoxHunt demonstrating Louvain clusters most strongly resemble regions of the developing pallium, which corresponds to the cerebral cortex. **e**, UMAP with individual cells annotated based on correlation to cell types from a published brain organoid atlas using SingleR. **f**, Heatmap of correlation scores for individual cells calculated using SingleR.

**Extended Data Fig. 3:**
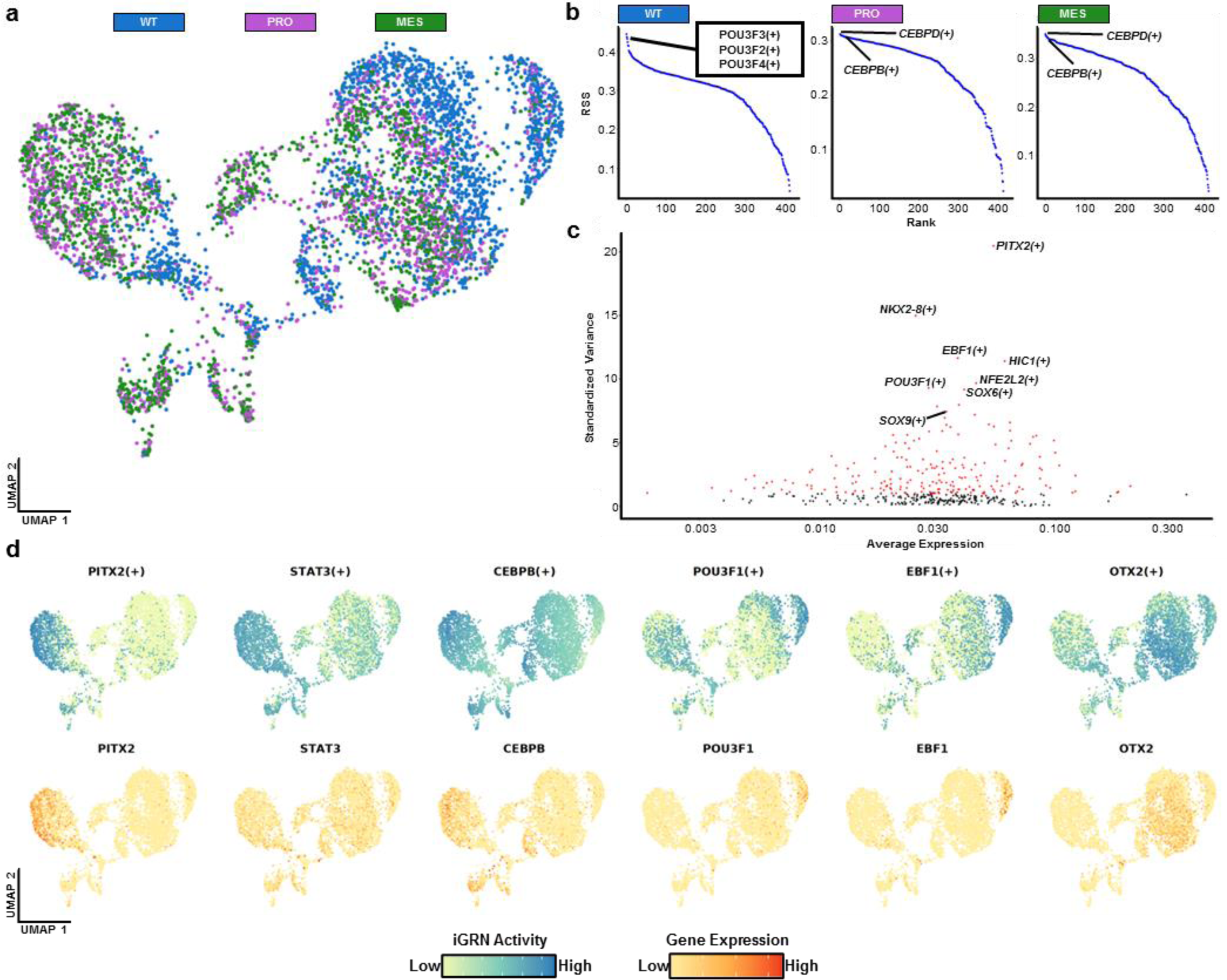
Analysis of iGRNs reveals activation of oncogenic gene regulatory networks in eGBOs. **a**, UMAP of WT, PRO, and MES organoids clustered by iGRN activity. **b**, Rank order plot of iGRN regulons sorted by regulon specificity score (RSS). **c**, Scatter plot of iGRN regulons. Red dots represent the top 50% of variable regulons based on standardized variance. **d**, Feature plots indicating iGRN activity of differentially active regulons (top) and gene expression of the corresponding TF (bottom).

**Extended Data Fig. 4:**
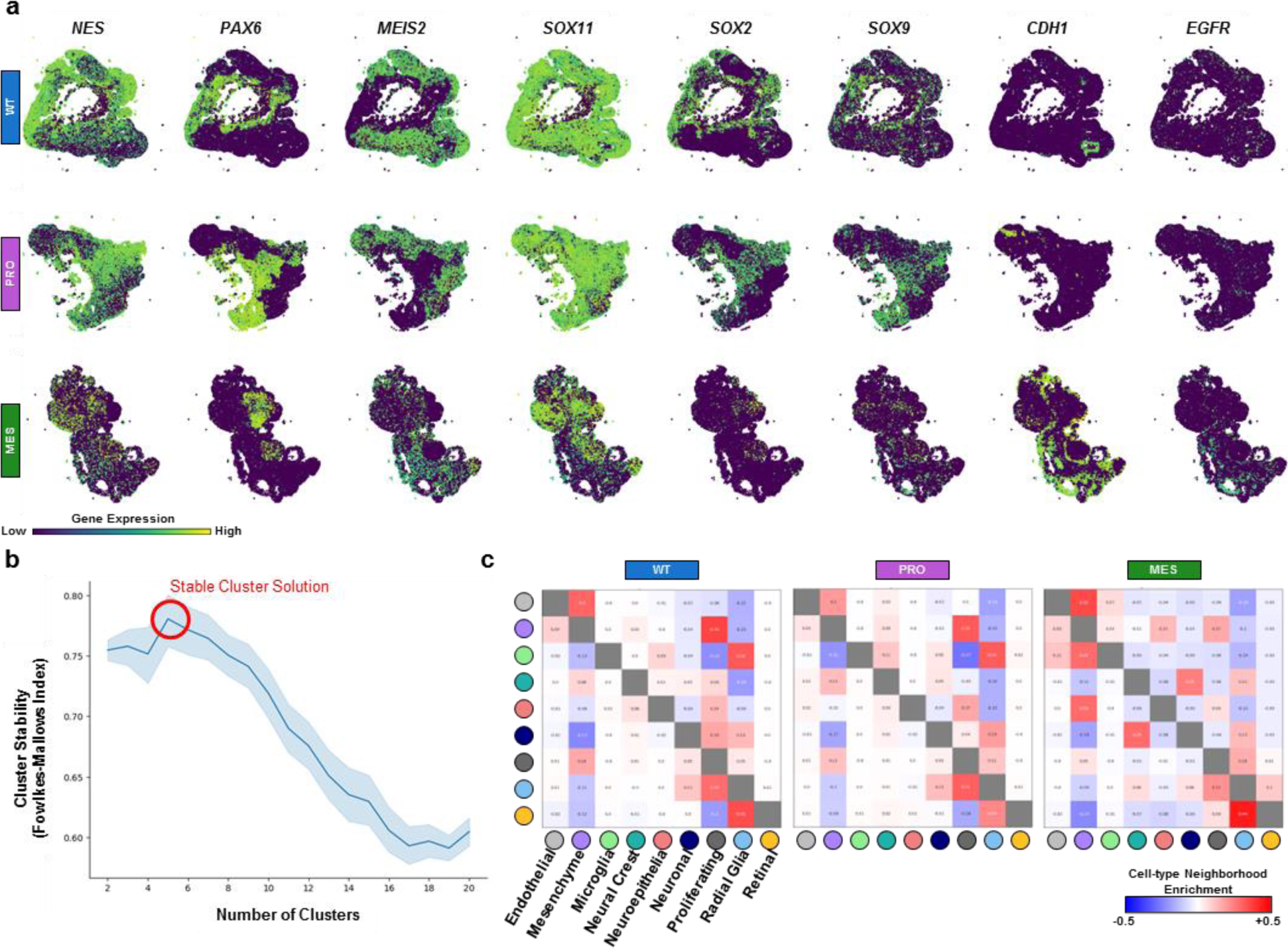
Analysis of spatially resolved transcriptomics in eGBOs. **a**, Spatial gene expression of genes in WT organoids and eGBOs. **b**, Cluster stability (y axis) for a range of number of clusters (x axis) for WT organoid and eGBO spatial sequencing samples. **c**, Cell type neighborhood enrichment heatmaps for WT organoids and eGBOs.

**Extended Data Fig. 5:**
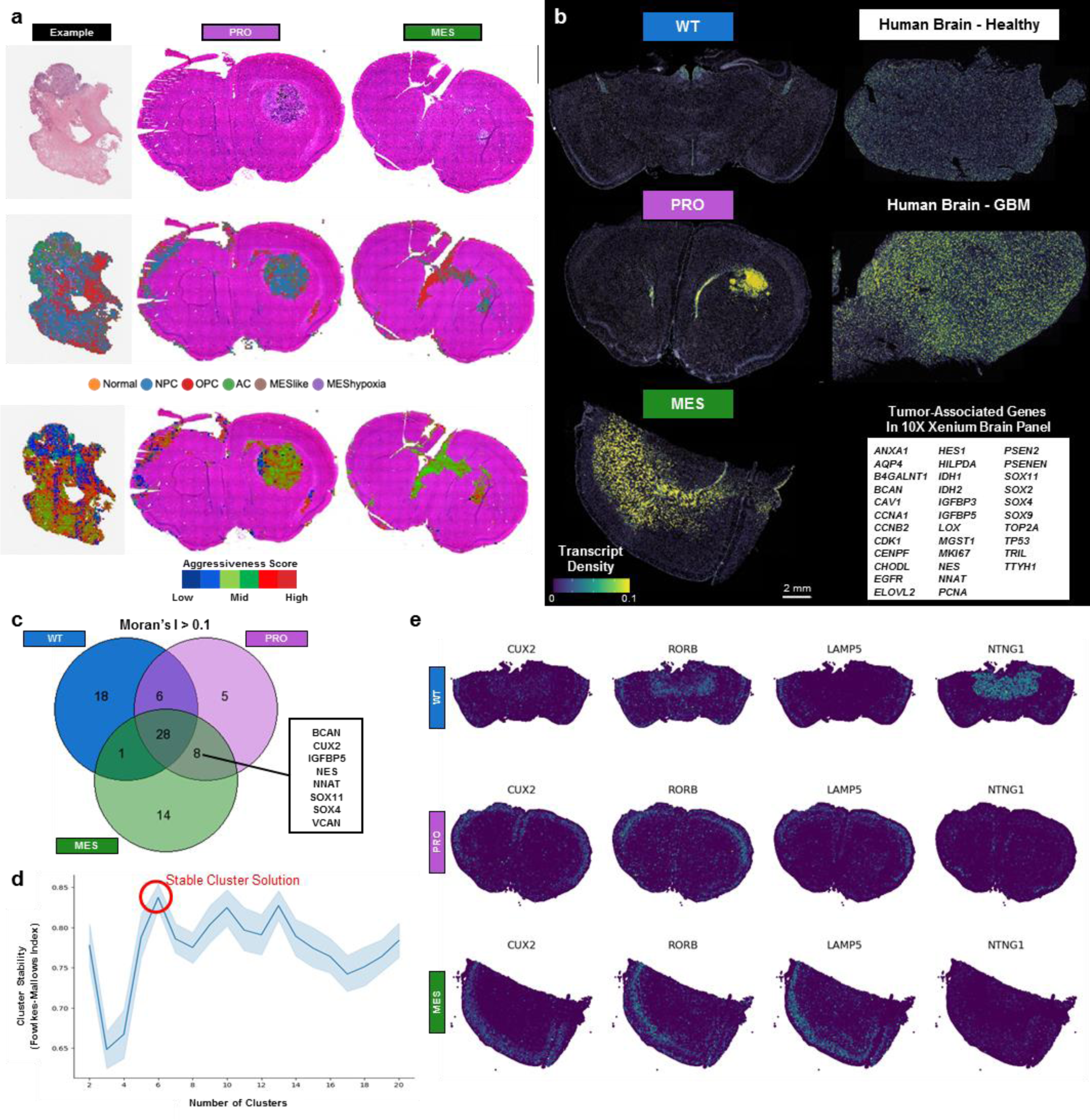
Spatially resolved transcriptomics analysis of tumors formed by eGBOs. **a**, H&E images (top row), predicted distribution of transcriptional subtypes (middle row), and predicted aggressive scores (bottom row) for a sample tumor from the TCGA cohort and tumors generated by eGBOs. **b**, Spatial gene expression profile of tumor-associated genes from the 10X Xenium brain panel. **c**, Venn diagram of spatially autocorrelated genes in brain sections from WT organoid and eGBO spatial sequencing samples. **d**, Cluster stability (y axis) for a range of number of clusters (x axis) for brain sections from WT organoid and eGBO spatial sequencing samples. **e**, Spatial gene expression of marker genes for different brain regions in spatial sequencing samples from WT organoid and eGBOs.

**Extended Data Fig. 6:**
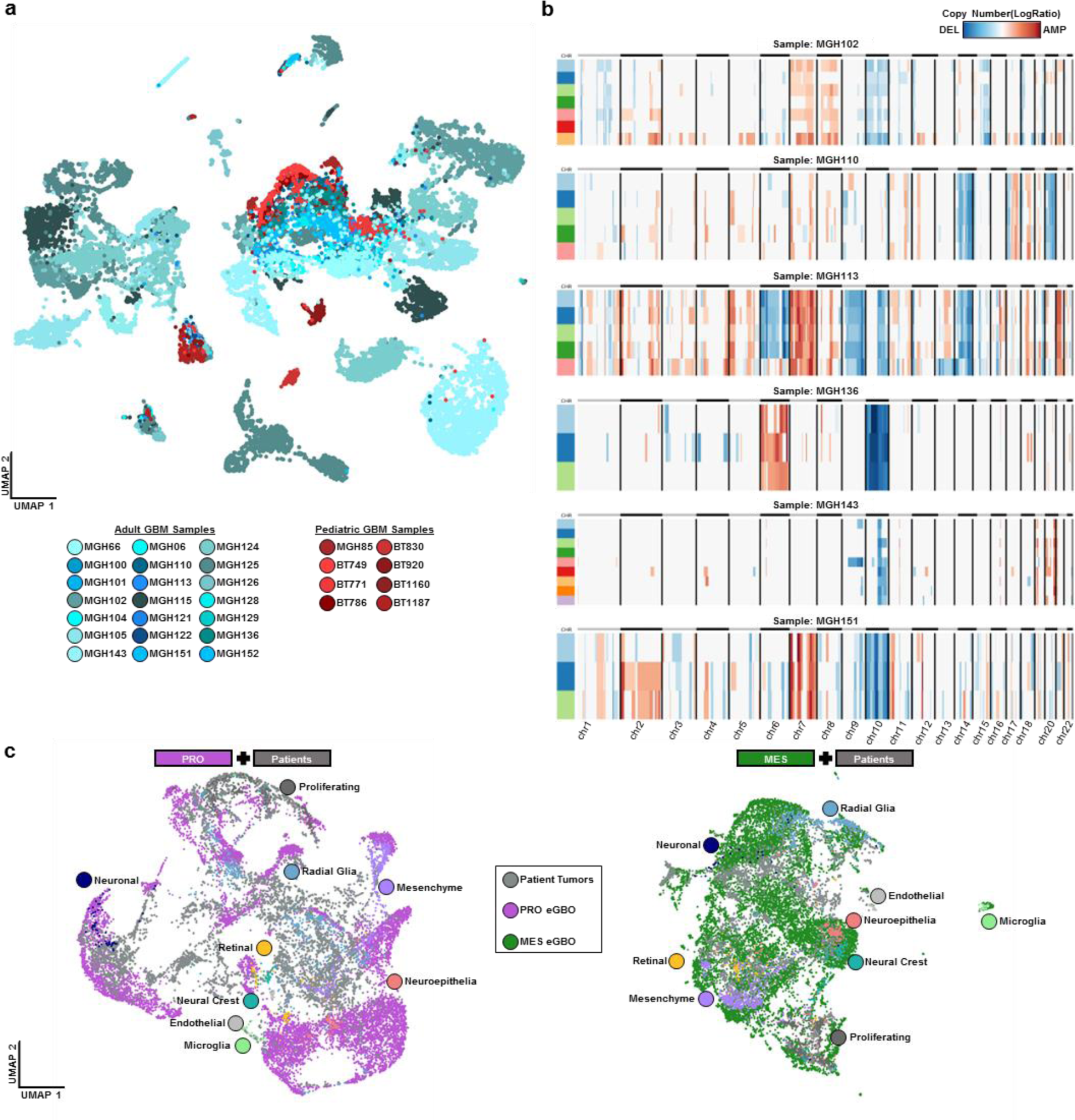
Inferred copy number aberrations in patient- and eGBO-derived tumors. **a**, UMAP of cells recovered from 29 GBM patient samples (21 adult and 8 pediatric). **b**, Compact representation of CNVs present in each subclone identified in a subset of patient GBM samples with similar mutations to eGBOs. **c**, UMAPs of integrated eGBO, eGBO-derived tumor, and patient-derived tumor cells indicating the distribution of eGBO cell types intermixed with tumor-derived cells.

